# Whole-genome sequencing of diverse 351 cultured prokaryotes including yet-unsequenced fastidious type strains

**DOI:** 10.1101/2025.05.25.655001

**Authors:** Shingo Kato, Sachiko Masuda, Arisa Shibata, Takashi Itoh, Mitsuo Sakamoto, Ken Shirasu, Moriya Ohkuma

**Affiliations:** Japan Collection of Microorganisms, RIKEN BioResource Research Center, Tsukuba, Japan; Plant Immunity Research Group, RIKEN Center for Sustainable Resource Science, Yokohama, Japan

## Abstract

Genome sequences provide fundamental information for both basic and applied life sciences. Whole-genome sequencing is now requested for describing novel prokaryotic species and designating their type strains, which serve as representative and well-characterized strains of the species. Indeed, the number of sequenced prokaryotic genomes has been rapidly increasing. However, a considerable number of isolated strains, particularly “fastidious” type strains such as strict anaerobes and slow growers, remain without genome sequence information. Here we report the whole-genome sequencing of 290 bacterial and 61 archaeal strains, including fastidious type strains, obtained from Japan Collection of Microorganisms (JCM) using a combination of short- and long-read sequencing technologies. The dataset includes 284 type strain genomes and 235 complete genomes. Notably, in the dataset, genomes of over 200 strains, including over 150 type strains, had not been made publicly available. Comparative genomic analysis suggests that some strains need to be assigned to novel taxa or reclassified. Functional gene survey indicates that some strains possess previously unrecognized potential for carbon fixation or bioactive secondary metabolite production. Our dataset will contribute to more accurate taxonomic classification, fill gaps in the phylogeny of prokaryotes, and provide insights into their physiology and ecology.

## INTRODUCTION

Whole-genome sequences serve as a fundamental blueprint for life, providing the foundation for a broad range of life sciences research. The genomes of prokaryotes, i.e., archaea and bacteria, are typically smaller (less than 10 Mbp) and simpler in structure, consisting of a single circular chromosome in most cases, than those of eukaryotes. The rapid development of DNA sequencing technologies and the increase in computational power have driven a significant expansion in the number of sequenced prokaryotic genomes over the years. In particular, type strains, which are representatives of species with well-characterized phenotypic information, have been a primary focus of large-scale whole-genome sequencing projects (1,2). In addition, since 2018, the determination of genome sequence of the type strain and comparison with those of related species have been required when describing novel prokaryotic species (3,4), resulting in a substantial accumulation of type strain genomes in public databases. Furthermore, recent advances in long-read sequencing technologies have enabled the re-sequencing and completion of type strain genomes (5,6). However, a considerable number of genomes from “fastidious” strains, such as slow-growing strict anaerobes with low maximum cell density, remain unsequenced due to the challenges associated with their cultivation and DNA extraction. Therefore, gaps in the prokaryotic phylogeny remain to be filled, and the full extent of their metabolic potentials remains unclear. Here, we report the whole-genome sequencing of 351 publicly available prokaryotic strains, including fastidious and/or previously unsequenced strains, obtained from the Japan Collection of Microorganisms (JCM), one of the international culture collections.

## MATERIAL AND METHODS

### Microbial strains and DNA extraction

We obtained 203 microbial strains from the Japan Collection of Microorganisms (JCM) at the RIKEN BioResource Research Center (BRC) in Tsukuba, Japan. These strains were cultivated under the growth conditions summarized in Table S1. Detailed information on the media used to culture each strain is available on the JCM website (https://jcm.brc.riken.jp/en/) and also deposited on the figshare website (https://doi.org/10.6084/m9.figshare.28756649.v1). Cells were harvested by centrifugation at 10,000 × *g* and stored at -80°C until DNA extraction. Genomic DNAs were extracted from the harvested cells using a DNeasy PowerLyzer Microbial Kit (Qiagen). Additionally, we obtained pre-extracted genomic DNAs for other 148 microbial strains from the DNA Bank of RIKEN BRC (Table S1). The purity and quantity of the DNA samples were assessed using a Qubit 3.0 Fluorometer (Thermo Fisher Scientific) with dsDNA quantification assay kits, as well as a Multiskan SkyHigh Microplate Spectrophotometer (Thermo Fisher Scientific) with a μDrop Plate (Thermo Fisher Scientific).

### Genome sequencing

We performed short-read sequencing on MiSeq or NextSeq 1000 instruments (Illumina). DNA libraries were constructed from aliquots of the DNA samples using the QIAseq FX DNA Library Kit (Qiagen) for sequencing on MiSeq and NextSeq instruments, and sequenced with the MiSeq Reagent Kit version 3 (600 cycles, 300 bp paired-end reads; Illumina) and the NextSeq 1000/2000 P1 XLEAP-SBS Reagent Kit (600 cycles, 300 bp paired-end reads; Illumina).

For long-read sequencing, we used a MinION device (Oxford Nanopore Technologies, ONT) and a PacBio Revio system (Pacific Biosciences). For ONT sequencing, DNA libraries were constructed using the Rapid Barcoding Kit 24 V14 (SQK-RBK114.24) and sequenced with R10.4.1 flow cells (FLO-MIN114). The raw data were basecalled using Dorado version 0.7.1 (https://github.com/nanoporetech/dorado) with the “sup” model. For PacBio sequencing, we first assessed the quality of the DNA samples using a Femto Pulse system (Agilent Technologies; Santa Clara, CA, United States). Subsequently, SMRTbell libraries were prepared using SMRTbell Express Template Prep Kit v2.0. On several libraries where necessary, size-selection was performed on the BluePippin system using a 0.75% agarose cassette (Sage Science; Beverly, MA, United States) with a 5-50 kb high-pass cutoff. The SMRTbell libraries were then bound to the sequencing polymerase enzyme using a Revio Polymerase kit. Shotgun genomic DNA sequence data were collected on SMRT Cells using HiFi sequencing protocols and Revio sequencing plate (PacBio). HiFi reads were generated using SMRTLink v13.1 with default parameters and extracted with a quality score >Q30.

### Genome assembly

The short reads were filtered using fastp version 0.23.4 (7) with the option “-l 200”. The long reads were filtered using Chopper version 0.7.0, implemented in NanoPack2 (8), with the options “-q 8 -l 1000”. In some cases, long read data exceeding the desired size were downsized to approximately 100× coverage using Filtlong (https://github.com/rrwick/Filtlong). The filtered long reads were assembled using Flye version 2.9.3 (9) or the Improved Phased Assembler version 1.8.0 (https://github.com/PacificBiosciences/pbipa) equipped with the Pacbio Revio system. If closed contigs for putative primary chromosomes were not obtained, hybrid assembly was performed using Unicycler version 0.5.0 (10) (short-read-first hybrid assembling) or Hybracter version 0.7.3 (11) (long-read-first hybrid assembling) with the short reads. For cases where only short reads were available, Unicycler was used for short-read-only assembly. The assembly software used for each strain is listed in Table S2. In this study, when the longest contig for the primary chromosome in the assembly was circularized, the genome was treated as a “complete” genome, with the exception of *Streptomyces* species, which may have linear chromosomes. For *Streptomyces* assemblies, the presence of terminal inverted repeats detected by BLASTn version 2.14.0 (12) was used to assess whether the linear genomes were complete.

### Genome analysis

The genome assemblies were then annotated using DFAST version 1.3.2 (13) with the option “--minimum_length 500 --use_prodigal --use_trnascan”. For additional annotation, we used METABOLIC version 4.0 (14) equipped with KEGG (15), HydDB (16), dbCAN2 (17) and MEROPS (18), eggNOG-mapper version 2.1.12 (19), and gapseq version 1.3.1 (20). Biosynthetic gene clusters (BGCs) were detected using antiSMASH version 7.1.0 (21). Plasmids and viruses/phages were identified using geNomad version 1.8.1 (22), mob-suite version 3.1.9 (23), and VirSorter version 2.2.4 (24).

The quality of the determined genomes was evaluated using CheckM version 1.2.3 (25) and CheckM2 version 1.0.2 (26). Taxonomic classification of strains based on the determined genomes was performed using GTDB-tk version 2.4.0 (27) with the reference database R226 (28). Average nucleotide identity (ANI) values among the determined genomes were calculated using FastANI version 1.34 (29). Average amino acid identity (AAI) values among the determined genomes were calculated using EzAAI version 1.2.3 (30). Values of the percentage of conserved proteins (POCP) were calculated using POCP-nf version 2.3.4 (31). Digital DNA-DNA hybridization (dDDH) values were determined using the Genome-to-Genome Distance Calculator (GGDC) version 3.0 (32). We used Type Strain Genome Server (32) and GTDB-tk to detect the closest publicly available genomes to determined ones, based on dDDH and ANI values, respectively. Alluvial diagrams were generated using RAWGraphs (https://www.rawgraphs.io). Plots and bar charts were generated using ggplot2 (https://ggplot2.tidyverse.org).

## RESULTS AND DISCUSSION

### Overview of the sequenced genomes

We determined the whole-genome sequences of 351 microbial strains, including 284 type strains (Fig. 1A; Table S2). Of the 351 genomes, 235 genomes (including 189 type strain genomes) were completed. The size of the determined genomes ranged from 1,265,941 bp for *Methanothermus sociabilis* JCM 10723^T^ to 10,631,713 bp for *Streptomyces phaeofaciens* JCM 4814^T^ (Fig. S1). The genome size of *M. sociabilis* JCM 10723^T^ is comparable to that of other small genomes of free-living organisms, such as ‘*Nitrosopelagicus brevis*’ CN25 (1.232 Mbp) and *Methanothermus fervidus* V24S^T^ (1.243 Mbp). The G+C content of the genomes varied widely, ranging from 24% for *Exilispira thermophila* JCM14728^T^ to 76% for *Cellulomonas pakistanensis* JCM18755^T^.

**Fig. 1.**
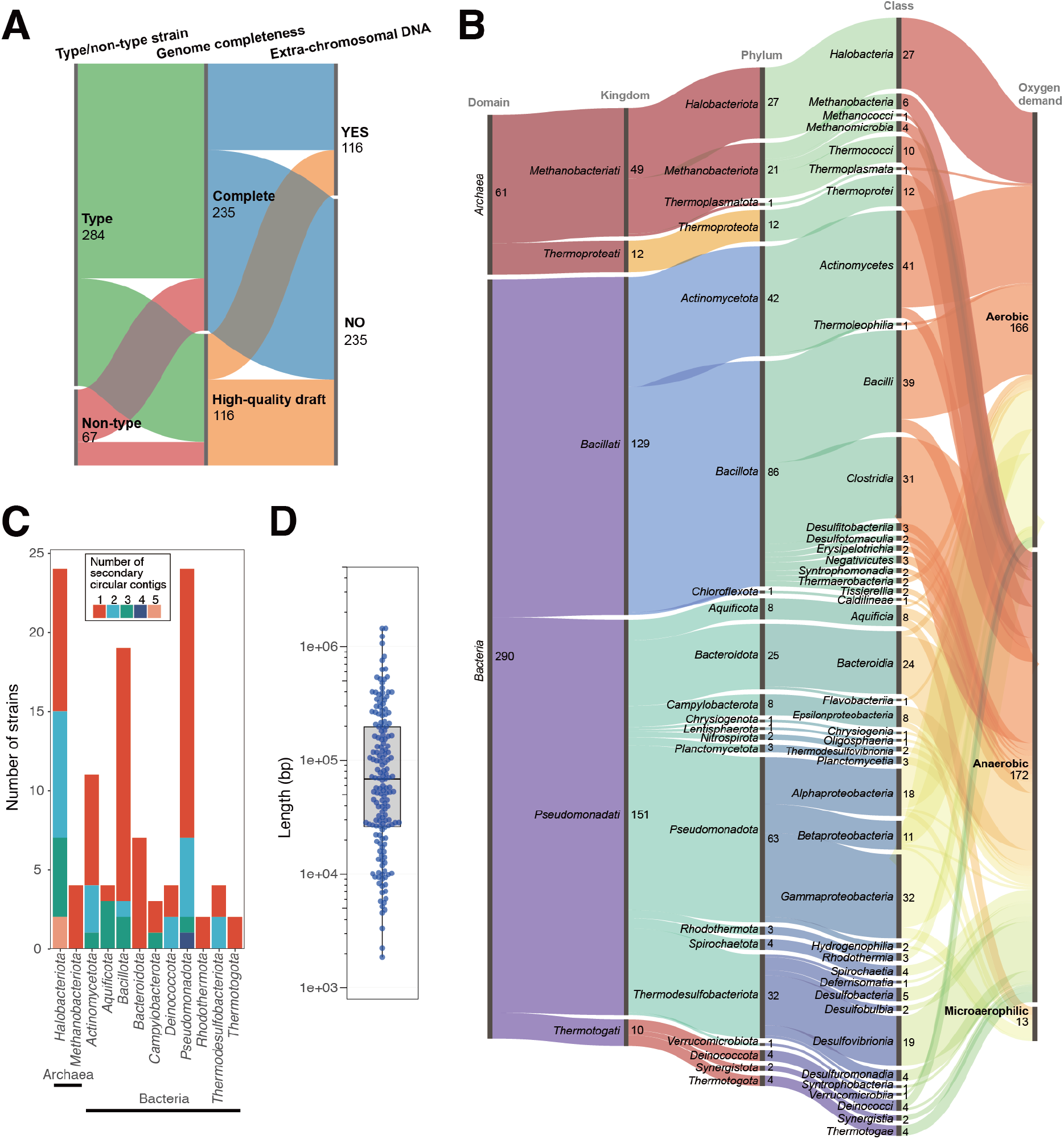
Overview of the determined genomes of the 351 strains. (**A**) Alluvial diagram showing the distribution of type and non-type strains, genome completeness, and the presence of extra-chromosomal DNA. (**B**) Alluvial diagram showing the taxonomic classification and oxygen requirement of the 351 strains. (**C**) Bar chart showing the number of strains harboring one or more secondary circular contigs. The number of secondary circular contigs is indicated by different colors as shown in the inset box. (**D**) Box plot showing the size distribution of the secondary circular contigs.

The dataset encompassed phylogenetically diverse prokaryotes, comprising 61 archaeal and 290 bacterial strains (Fig. 1B; Table S3). The taxonomic classification, based on the International Code of Nomenclature of Prokaryotes (ICNP), showed that our dataset spanned 4 archaeal phyla with 7 classes, and 18 bacterial phyla with 36 classes. In addition, these genomes represented physiologically diverse microorganisms, including anaerobes and microaerophiles. Overall, our dataset covers a broad range of prokaryotic taxa, despite being much smaller number than the total of publicly-available genomes.

Although some genomes showed high values of contamination level, many of these genomes were completed (Table S2). For instance, the determined genome of *“Desulfosporomusa polytropa”* JCM 32836^T^ was fully completed, comprising two circular contigs of 5,636,488 bp and 261,751 bp, but showed contamination values of up to 12.1%. Notably, the strain heterogeneity value was zero, indicating no actual contamination.

Therefore, we considered that the high values of contamination level observed in this strain and also others were not due to actual contamination, but rather a limitation of the checking ability for diverse taxa in the software used in this study. Indeed, no 16S rRNA gene sequences from potential contaminants were detected in any of the assemblies. Besides, some complete genomes showed low values of completeness level, *e*.*g*., <95% for *Oligosphaera ethanolica* JCM 17152^T^, which can also be attributed to the inadequacy of the software. The high-quality genome data obtained from pure cultures in this study provide a robust basis for more accurate assessments of genome completeness and contamination levels.

The genome sequencing of the isolated strains indicated the presence of extra-chromosomal DNA fragments, such as secondary replicons (33) including plasmids and chromids, or phages/viruses, in addition to primary chromosomes. Metagenomics of complex microbial communities in natural environments could detect such extra-chromosomal DNA fragments, but often fail to identify their host organisms. In this study, we found that 76 out of 235 complete genomes contained two or more contigs (Table S2), suggesting the presence of extra-chromosomal DNA fragments. In particular, these secondary contigs in 68 of the 76 genomes included circular contigs. Furthermore, 40 draft genomes contained smaller circular contigs than each of the longest contig, which could also represent extra-chromosomal DNA fragments. The circular contigs (up to 5 per genome) of extra-chromosomal DNAs were found in the genomes in 12 of the 22 phyla (Fig. 1C). The size of these circular contigs ranged from 1,847 to 1,436,886 bp (Fig. 1D), which were likely to include megaplasmids, chromids, or secondary chromosomes. Remarkably, 35 of the circular contigs were not detected as plasmids or phages/viruses using existing tools, suggesting that these were novel types of secondary replicons or phages/viruses. The results will contribute to expand the knowledge of extra-chromosomal DNAs of prokaryotes.

### Potential proposal of novel taxa and reclassification

Notably, 209 (38 archaeal and 171 bacterial) of the 351 genomes, including 166 type strain genomes, were being released from public databases for the first time (Fig. 2; Table S2). Of the 166 type strain genomes, the genomes of 78 strains, *e*.*g*., *Hydrogenobaculum acidophilum* JCM 8795^T^, showed low similarity (<95% ANI or <70% dDDH, commonly used thresholds for species-level definition (34,35)) to any of publicly available genomes derived from isolates or metagenome-assembled genomes (MAGs) (Table S3). This indicates that these genomes are the first to be publicly released at the species level. Other 17 genomes showed high similarity (>95% ANI) to previously-reported MAGs or genomes of yet-unnamed isolates. For instance, the genome of *Methanothermobacter crinale* JCM 17393^T^ showed high similarity (99.6% ANI) to the MAG 41_258 (accession no. GCA_001507955.1) recovered from an oil reservoir (36). It should be noted that some of the counts of genomes reported in the present study may be subject to change due to ongoing updates of public databases.

**Fig. 2.**
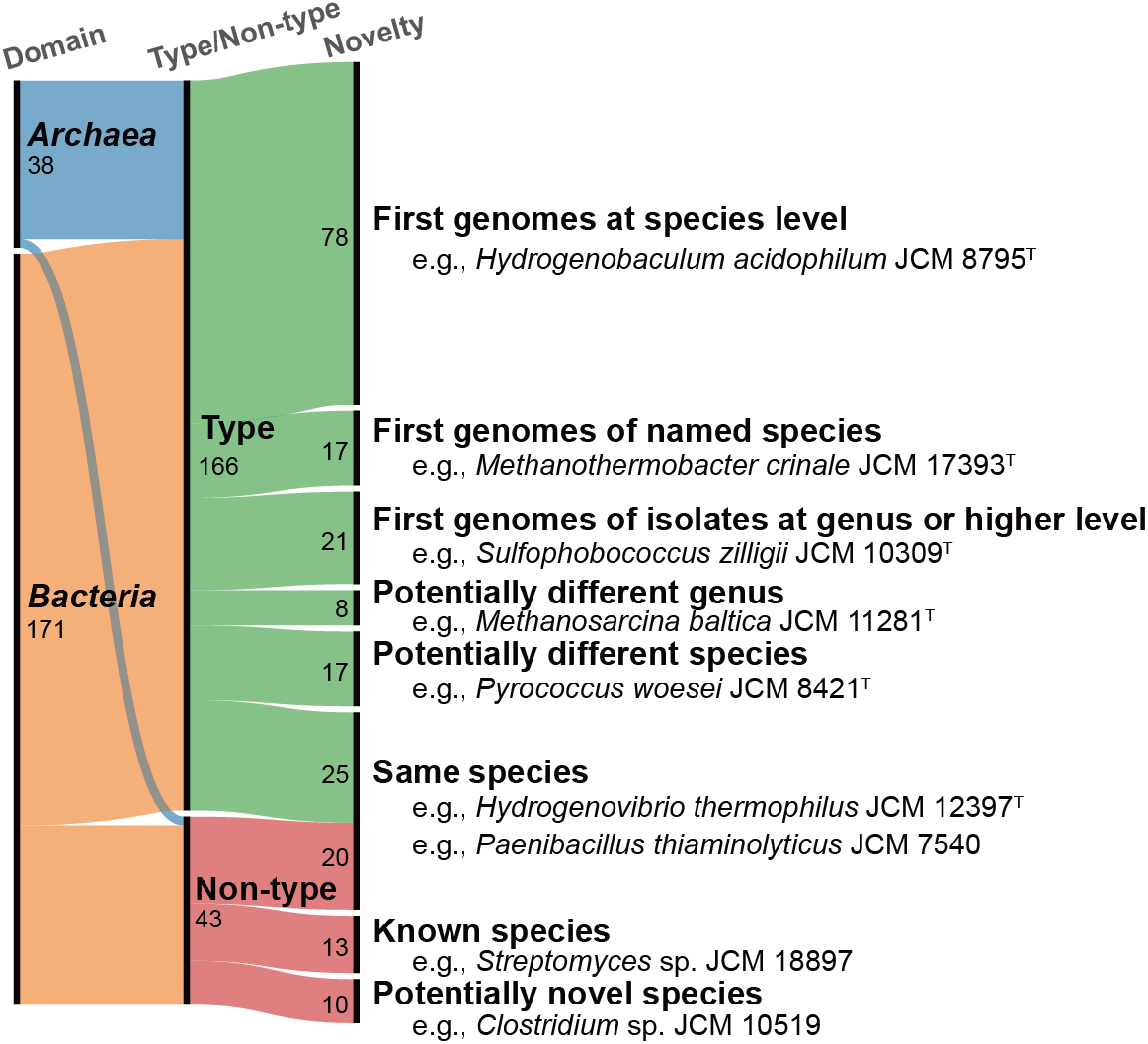
Genomic and taxonomic novelty. Alluvial diagram showing the genomic and taxonomic novelty of the 209 strains with newly released genomes in this study. Details are provided in the main text and in Table S3.

Of the 166 newly released type strain genomes, other 21 genomes were the first type strain genomes to be publicly released at the genus level, based on the GTDB-based taxonomic classification. For instance, the genome of *Sulfophobococcus zilligii* JCM 10309^T^ did not match any genomes of isolates, but showed high similarity (98.7% ANI) to a MAG (UBA285, accession no. GCA_002495025.1), which was assigned to the genus-level clade “g_UBA285” within the family-level clade “f_Desulfurococcaceae”. This result indicates that the genus-level clade “g_UBA285” corresponds to the genus *Sulfophobococcus*.

Similarly, we released the first type strain genomes for other 20 validly named genera, including *Stetteria* and *Thermodiscus* in the domain *Archaea*, as well as *Vulcanithermus* and *Thioreductor* in the domain *Bacteria*. Moreover, some genomes of the 21 type strains had a potential for the assignment to novel taxa at the family or higher level. According to the GTDB-based classification (Table S3), our genome dataset potentially encompassed 1 novel class, 3 novel orders, and 4 novel families. For instance, *Exilispira thermophila* JCM 14728^T^, a member of the phylum *Spirochaetota*, could represent a novel family, order, or even class. Similarly, *Endothiovibrio diazotrophicus* JCM 17961^T^ belonging to the class *Gammaproteobacteria* could represent a novel family and order. *Thioprofundum lithotrophicum* JCM 14586^T^, belonging to the family *Thioprofundaceae*, could represent a novel order. Both *Brassicibacter thermophilus* JCM 30480^T^ and *Thiofractor thiocaminus* JCM 15747^T^ could represent a novel family. Further phylogenetic analysis is needed to support the assignment to these novel taxa.

The GTDB-based classification suggested that other 8 type strain genomes, currently assigned to already known genera, could represent novel genera (Fig. 2). For example, *Methanosarcina baltica* JCM 11281^T^ was classified as “g_JAQVBP01”, which is distinct from “g_Methanosarcina” and the corresponding genus *Methanosarcina*. This result is consistent with a previous study reporting that *M. baltica* can be physiologically and phylogenetically distinguished from other species in *Methanosarcina* (37). Among species in *Methanosarcina*, the most similar type strain to *M. baltica* JCM 11281^T^ was *Methanosarcina subterranea* JCM 15540^T^, whose genome was determined in this study and classified as “g_Methanosarcina”. The comparisons between these two strains showed the values of 94.9% (16S rRNA gene similarity), 72.9% (AAI), and 61.9% (POCP). Given the reported values for genus boundary (90-99% for 16S rRNA gene similarity, 65–72% for AAI, and 50– 60% for POCP) (38-41), *M. baltica* JCM 11281^T^ is standing on the boundary of the genus threshold. These results imply that *M. baltica* JCM 11281^T^ belongs to a distinct genus from *Methanosarcina*, although further careful analysis is needed to conclude this notion.

Other 17 type strain genomes showed high similarity (≥95% ANI or ≥70% dDDH) to those of different species (Fig. 2), suggesting that taxonomic reclassification should be considered. For example, the determined genome of *Pyrococcus woesei* JCM 8421^T^ showed high similarity to that of *Pyrococcus furiosus* DSM 3638^T^ (accession no. GCA_000007305.1), with ANI and dDDH values of 99.6% and 95.2%, respectively. This result supports the notion that these two strains belong to the same species, *i*.*e*., *Pyrococcus furiosus*, which has been previously noted due to the high similarities on their gene sequences and physiological characteristics (42). Similarly, the genome of *‘Thermococcus marinus’* JCM 11825^T^, a proposed novel species with an as-yet unvalidated name (43), showed high similarity to that of *Thermococcus eurythermalis* A501^T^ (accession no. GCA_000769655.1), with ANI and dDDH values of 99.2% and 90.5%, respectively. Our dataset includes 14 additional strains that have been proposed to represent novel species, but whose scientific names have not yet been validated (Table S3). Given the strong recommendation for whole-genome sequences in the field of prokaryotic taxonomy since 2018, our dataset is expected to facilitate taxonomic reclassification and assessment of the validity of proposed scientific names.

Among the newly released 43 genomes of non-type strains, 10 genomes showed low similarity to any publicly available or determined genomes of type strains, indicating that they represented probable novel species. For instance, the genome of *Clostridium* sp. JCM 10519 (=NkU-1) showed a low similarity to its closest relative, *Lacrimispora saccharolytica* (formerly *Clostridium saccharolytica*) strain WM1^T^ (accession no. GCA_000144625.1), with ANI and dDDH values of 85.0% and 29.8%, respectively. Indeed, despite a high 16S rRNA gene similarity of 98.9% between the two strains, their physiological characteristics are distinct as reported previously (44). Further analyses of *Clostridium* sp. JCM 10519 and the other 9 strains are expected to provide additional supporting evidence for describing novel species. Other 13 genomes of unidentified species showed high similarities (≥95% ANI or ≥70% dDDH) to those of known species. For instance, the genome of *Streptomyces* sp. JCM 18897 was highly similar to that of *Streptomyces albidoflavus* NRRL B-1271^T^ with 99.0% ANI and 95.9% dDDH, and thus *Streptomyces* sp. JCM 18897 could be assigned to the species *Streptomyces albidoflavus*.

The remained 25 genomes of type strains showed high similarities to the genomes of the same species of non-type strains. For instance, the determined genome of *Hydrogenovibrio thermophilus* JCM 12397^T^ showed a high similarity (97.1% ANI) to the genome of *H. thermophilus* JR-2 (a non-type strain; accession no. GCF_004028275.1) recovered from a deep-sea hydrothermal vent field (45). The remained 20 genomes of non-type strains, e.g., *Paenibacillus thiaminolyticus* JCM 7540, showed high similarities to the previously reported genomes of the same species of type- or non-type strains.

### Potential for previously unrecognized metabolism

Genome analysis of isolates not only enables the prediction of previously unrecognized metabolic processes by identifying known genes but also facilitates the discovery of novel pathways underlying experimentally validated metabolisms. In the determined genomes, we identified genes associated with a variety of known metabolic functions (Table S4). Overall, the gene context is largely consistent with reported metabolic activities characterized through cultivation. For example, key genes for methanogenesis (*mcr* for methyl-coenzyme M reductase) and methane oxidation (*pmo* for particulate methane monooxygenase) were detected in the genomes of methanogenic archaea belonging to *Methanobacteriota*, such as *Methanobacterium movens* JCM 15415^T^, and methane-oxidizing bacteria belonging to *Pseudomonadota*, such as *Methylosoma difficile* JCM 14076^T^, respectively (Table S4).

However, several predicted metabolic capabilities remain unverified. In this study, we further focused on the potential for carbon fixation and secondary metabolite production, both of which are critical for applications in carbon neutrality and medical and agricultural sciences, directly relevant to human activities.

#### Potential for carbon fixation

To date, seven pathways are known to be involved in autotrophic carbon fixation (46,47), including the Calvin-Benson-Bassham (CBB) cycle, the reverse tricarboxylic acid (rTCA) cycle, and the Wood–Ljungdahl (WL) pathway.

Additionally, some autotrophs can fix CO_2_ via the reversed oxidative TCA (roTCA) cycle, which does not require key enzymes such as citryl-CoA synthetase (CCS) or ATP-citric lyase (ACL) in the rTCA cycle, but is driven only by the enzymes used in the “normal” oxidative TCA cycle (48,49) with a key enzyme of ferredoxin-dependent 2-oxoglutarate synthase (50).

We showed that the 67 genomes possessed the complete or nearly complete gene set with the key genes for the CBB cycle, rTCA cycle, WL pathway, and roTCA cycle (Fig. 3A; Table S5). As expected, most of the 67 strains have been reported to be autotrophs. However, we also found the key genes in the genomes of some strains that have not been reported as autotrophs. For example, *Mycolicibacterium crocinum* JCM 16369^T^ and *Mycolicibacterium pallens* JCM 16370^T^ have not been tested for the capability of carbon fixation (51), whereas their complete genomes possessed the complete gene set for the CBB cycle. Indeed, some autotrophic species in the genus *Mycolicibacterium* have been reported (52), suggesting that the two strains may also be capable of autotrophic growth.

**Fig. 3.**
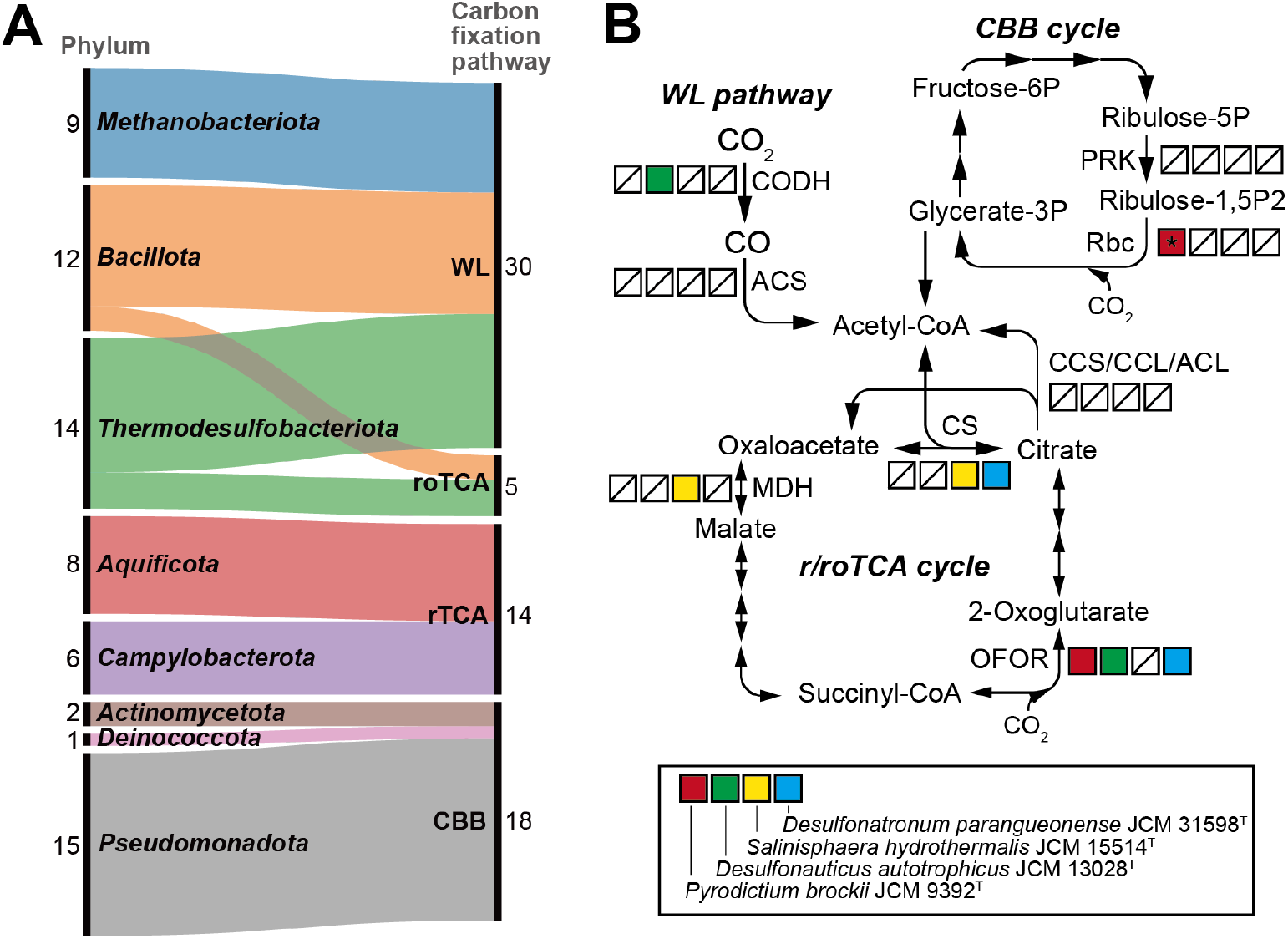
Carbon fixation pathways encoded in the genomes. (**A**) Alluvial diagram showing the phylum-level classification of the 67 strains possessing key genes for each carbon fixation pathway. WL, the Wood–Ljungdahl pathway; roTCA, the reversed oxidative tricarboxylic acid cycle; rTCA, the reverse tricarboxylic acid cycle; CBB, the Calvin-Benson-Bassham cycle. (**B**) Presence of key genes for each carbon fixation pathway in the genomes of the four autotrophic strains highlighted in the box. *A gene for RuBisCO form III, but not forms I or II, was detected in the genome of *Pyrodictium brockii* JCM 9392^T^ (see the main text for details). CCS, citryl-CoA synthetase; CCL, citryl-CoA lyase; ACL, ATP-citrate lyase; CODH, carbon-monoxide dehydrogenase; ACS, acetyl-CoA synthase; PRK, phosphoribulokinase; Rbc, ribulose-bisphosphate carboxylase; CS, citrate synthase; OFOR, 2-oxoacid:ferredoxin oxidoreductases; MDH, malate dehydrogenase.

Notably, as shown in Fig. 3B, the key genes for the seven carbon fixation pathways and even roTCA cycle were incompletely found in the genomes of the four strains, *i*.*e*., *Pyrodictium brockii* JCM 9392^T^ (53), *Desulfonauticus autotrophicus* JCM 13028^T^ (54), *Desulfonatronum parangueonense* JCM 31598^T^ (55), and *Salinisphaera hydrothermalis* JCM 15514^T^ (56), all of which have been reported as autotrophs. Except for JCM 31598^T^, complete genomes of the three strains were determined in this study. Therefore, they potentially fix CO_2_ via unknown carbon fixation pathways. In the case of *S. hydrothermalis* JCM 15514^T^, the gene for ribulose-1,5-bisphosphate carboxylase/oxygenase (RuBisCO), a key enzyme of the CBB cycle, has been detected by PCR cloning-sequencing (56). However, neither RuBisCO gene nor other key genes for carbon fixation were found in the determined complete genome sequence nor in the previously reported draft genome sequence (accession no. APNE00000000). In the complete genome of *P. brockii* JCM 9392^T^, we found a gene for RuBisCO form III, which may not involve in carbon fixation (57), and no gene for phosphoribulokinase (PRK), another key enzyme of the CBB cycle. For the above four strains, further analyses including re-evaluation of their autotrophy will be needed to reveal the presence of novel carbon fixation pathways.

#### Potential for secondary metabolite production

Bioactive secondary metabolites, including antibiotics, are crucial targets for applications in medicine and agriculture, as well as for understanding interactions among organisms in ecology. The genes responsible for producing secondary metabolites are often encoded in biosynthetic gene clusters (BGCs), which have been found in the genomes of diverse bacteria and archaea (58-60).

We identified 1,696 BGCs in 281 of the determined 351 genomes (Fig. 4; Table S6), not only from aerobic bacteria including well-studied *Streptomyces* species of the phylum *Actinomycetota*, but also from archaea and anaerobic bacteria, which have been less studied (61). Of the 235 complete genomes, 222 (94.9%) possessed one or more BGCs. The BGC counts were roughly correlated with the genome sizes in our dataset (r^2^ = 0.524) (Fig. 4A), a trend that has been previously reported for *Actinomycetota* genomes (62). The types of BGCs, such as non-ribosomal peptide synthetases (NRPSs), polyketide synthases (PKSs), and ribosomally synthesized and post-translationally modified peptides (RiPPs), varied among taxa (Fig. 4B). For instance, PKS-categorized BGCs were only found in the bacterial genomes. Terpene-categorized BGCs were rare in the genomes of *Methanobacteriota* and *Thermodesulfobacteriota*, both of which include obligate anaerobes. In contrast, RiPP-categorized BGCs were widely detected among the bacterial and archaeal phyla.

**Fig. 4.**
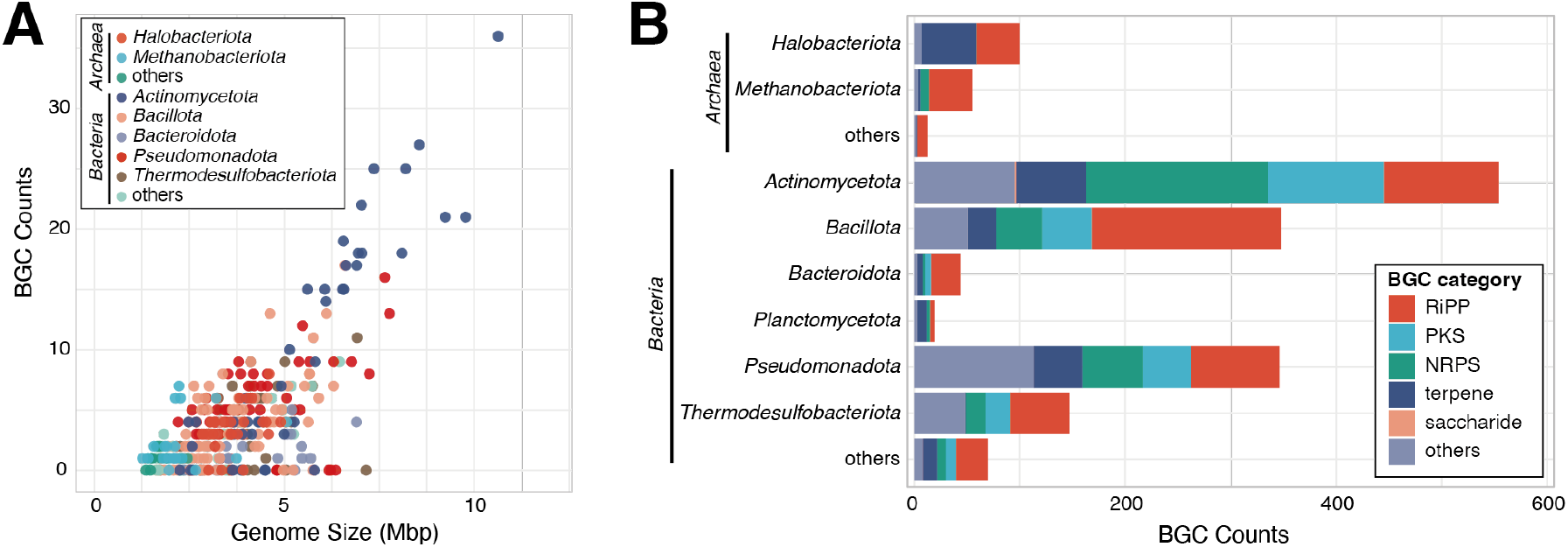
Biosynthetic gene clusters (BGCs) encoded in the genomes. (**A**) Plot showing the relationship between genome size and the number of BGCs. (**B**) Bar chart showing the number of BGCs for each category. RiPP, ribosomally synthesized and post-translationally modified peptide; PKS, polyketide synthase; NRPS, non-ribosomal peptide synthetase.

Over 20 BGCs per genome were exclusively detected from the phylum *Actinomycetota*, with the highest number (36 BGCs) found in the genome of *Streptomyces phaeofaciens* JCM 4814^T^, although some *Streptomyces* species have been reported to harbor over 70 BGCs per genome (62,63). Notably, 10 or more BGCs per genome were detected from the phyla *Bacillota, Pseudomonadota*, and *Thermodesulfobacteriota*, in addition to *Actinomycetota*. In obligate anaerobic bacteria, up to 11 BGCs were found in the genomes of *Desulfoconvexum algidum* JCM 16085^T^ (*Thermodesulfobacteriota*) and *Clostridium nitrophenolicum* JCM 14030^T^ (*Bacillota*), whose genomes were first determined and completed in this study. In contrast, archaeal genomes had fewer BGCs, which correlated with their smaller genome sizes (Fig. 4A). The highest number (8 BGCs) was found in the aerobic halophile *Halorubellus litoreus* JCM 17117^T^, followed by 7 BGCs in the obligate anaerobic methanogen *Methanobacterium movens* JCM 15415^T^. Although the functions of most secondary metabolites produced by BGCs remain unknown, the determined genomes of the isolated strains, especially those of not-well-studied archaea and anaerobic bacteria, will serve as a valuable foundation for genome-based mining of novel bioactive compounds.

## Conclusion

In this study, we determined the whole-genome sequences of 351 prokaryotic strains, spanning a broad range of archaeal and bacterial taxa, including previously unsequenced “fastidious” type strains. The dataset of complete or near-complete genomes provides a robust basis for more accurate assessments of genome completeness and contamination, for extending our knowledge of prokaryotic extra-chromosomal DNAs, and for describing novel taxa and reclassifications. Furthermore, it also enables the prediction of previously unrecognized metabolic processes and the discovery of novel pathways underlying experimentally validated metabolisms. Importantly, the genome dataset was constructed from isolated strains publicly available from culture collections, and therefore, genome-driven hypotheses can be verified by cultivation experiments with the isolates.

## ACKNOWLEDGEMENTS

We would thank Hiromi Omokawa, Nahomi Noda, Kai Zhang, Naomi Sakurai, Nagisa Sato, Michiru Shimizu, and Koji Suzu for their technical assistance.

## AUTHOR CONTRIBUTIONS

Shingo Kato: Conceptualization, Formal analysis, Methodology, Validation, Writing— original draft, review & editing. Sachiko Masuda, Arisa Shibata, Takashi Itoh, and Mitsuo Sakamoto: Formal analysis, Methodology, Writing—review & editing. Ken Shirasu and Moriya Ohkuma: Conceptualization, Validation, Writing—review & editing.

## SUPPLEMENTARY DATA

Supplementary Data are available at journal online.

## DATA AVAILABILITY

Sequence data of raw reads and genome assemblies have been deposited in GenBank/DDBJ/EMBL under the BioProject numbers, PRJDB20344, PRJDB20346 and PRJDB20693, PRJDB20694, respectively. Supplementary data are available in the figshare website (https://figshare.com/s/b650f6ad54b2e15244fd).

