## Supplementary Data for "Whole-genome sequencing of diverse 351 cultured prokaryotes including yet-unsequenced fastidious type strains"

**Supplementary tables**

Table S1. List of the 351 strains used in this study.
Table S2. Genomic features of the 351 strains.
Table S3. Taxonomic classification of the 351 strains.
Table S4. Gene context in the 351 genomes determined in this study.
Table S5. List of autotrophic and potentially autotrophic 67 strains.
Table S6. Biosynthetic gene clusters (BGCs) detected in 281 genomes.

**Supplementary figure**

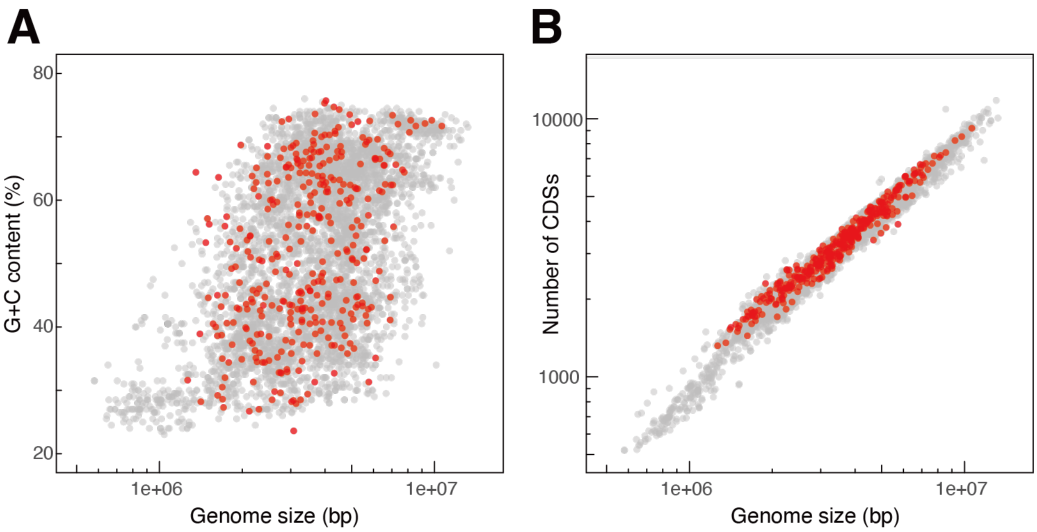


**Fig. S1. Genome size, G+C content, and the number of protein-coding regions (CDSs).** Plots showing the relationship between genome size and (**A**) G+C content or (**B**) the number of CDSs. Points represent the 351 genomes determined in this study (red) and complete prokaryotic genomes collected from NCBI RefSeq (gray).
